# A viral ring nuclease anti-CRISPR subverts type III CRISPR immunity

**DOI:** 10.1101/778746

**Authors:** Januka S Athukoralage, Stephen McMahon, Changyi Zhang, Sabine Grüschow, Shirley Graham, Mart Krupovic, Rachel J Whitaker, Tracey Gloster, Malcolm F White

## Abstract

The CRISPR system provides adaptive immunity against mobile genetic elements in bacteria and archaea. On detection of viral RNA, type III CRISPR systems generate a cyclic oligoadenylate (cOA) second messenger^1–3^, activating defence enzymes and sculpting a powerful antiviral response that can drive viruses to extinction^4,5^. Cyclic nucleotides are increasingly implicated as playing an important role in host-pathogen interactions^6,7^. Here, we identify a widespread new family of viral anti-CRISPR (Acr) enzymes that rapidly degrade cyclic tetra-adenylate (cA_4_). The viral ring nuclease (AcrIII-1) is the first Acr described for type III CRISPR systems and is widely distributed in archaeal and bacterial viruses, and proviruses. The enzyme uses a novel fold to bind cA_4_ specifically and utilizes a conserved active site to rapidly cleave the signalling molecule, allowing viruses to neutralise the type III CRISPR defence system. The AcrIII-1 family has a broad host range as it targets cA_4_ signalling molecules rather than specific CRISPR effector proteins. This study highlights the crucial role of cyclic nucleotide signalling in the conflict between viruses and their hosts.

---

Type III CRISPR-Cas systems synthesise the signalling molecule cyclic oligoadenylate (cOA) from ATP^1,2^ when they detect viral RNA. cOA molecules are synthesised with a range of ring sizes with 3-6 AMP subunits (denoted cA_3_, cA_4_ etc.) by the cyclase domain of the Cas10 protein^1–3,9,10^. cOA binds to a specific protein domain, known as a CARF (CRISPR Associated Rossman Fold) domain. CARF domains are found fused to a variety of effector domains that are known or predicted to cleave RNA, DNA, or function as transcription factors^11^. The best characterised CARF protein family is the Csx1/Csm6 family of HEPN (Higher Eukaryotes and Prokaryotes, Nucleotide binding) ribonucleases, which are activated by cOA binding and cleave RNA with minimal sequence dependence^1–3^. A number of studies have demonstrated that the cOA signalling component of type III systems is crucial for effective immunity against viruses^4,12–15^, highlighting the importance of this facet of CRISPR immunity.

Recently, we identified a cellular enzyme in *Sulfolobus solfataricus*, hereafter referred to as the Crn1 family (for “CRISPR associated ring nuclease 1”), that degrades cA_4_ molecules and thus deactivates the Csx1 ribonuclease *in vitro*^16^. These enzymes exhibit very slow kinetics, and are thought to act by mopping up cA_4_ molecules in the cell without compromising the immunity provided by the type III CRISPR system. Unsurprisingly, viruses have responded to the threat of the CRISPR system by evolving a range of anti-CRISPR (Acr) proteins, which are used to inhibit and overcome the cell’s CRISPR defences (reviewed in^17^). Acr’s have been identified for the type I-D^18^, I-F, II-A and V-A effector complexes (reviewed in^17,19,20^), numbering over 40 families^21^, but importantly not for type III systems. We focussed on one of the protein families, DUF1874, conserved and widespread in a variety of archaeal viruses and plasmids but also bacteriophages and prophages (Extended data figure 1) and has no known function. Structures are available for several family members, including gp29 of *Sulfolobus islandicus* rod-shaped virus 1 (SIRV1)^22^ and B116 of *Sulfolobus* turreted icosahedral virus (STIV)^23^, which is expressed early in the STIV infection cycle^24^ and is important for normal virus replication kinetics^25^.

Given the widespread distribution and important role in viral infection, we explored the possibility of an Acr function for this viral protein family. To investigate this, we deleted the genes for the type I-A system in *Sulfolobus islandicus* M.16.4 carrying *pyrEF* and *argD* deletions^26^ so that it had only a type III-B CRISPR system for defence^27^. We challenged this strain with the archaeal virus SSeV (Figure. 1a), a lytic virus isolated from Kamchatka Russia with an exact CRISPR-spacer match of 100% in M.16.4 and several other potentially active CRISPR-spacers (Rowland et al. *in prep*). SSeV lacks a *duf1874* gene and fails to form plaques on a lawn of *S. islandicus* M.16.4 with type III-B CRISPR defence (Figure 1a). However, the same cells expressing the SIRV1 gp29 gene from a plasmid are readily infected, giving rise to plaque formation. These data suggest that SIRV1 gp29 is functioning as an anti-CRISPR specific for type III CRISPR defence.

**Figure 1.**
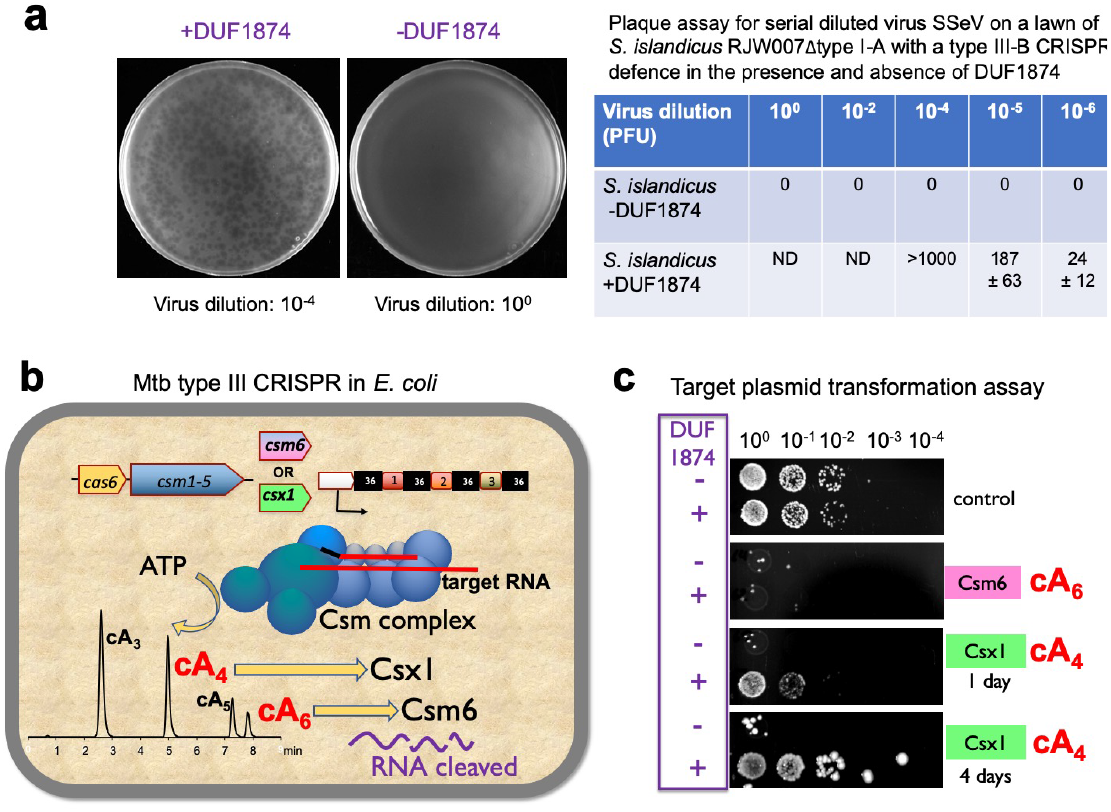
DUF1874 is an anti-CRISPR specific for cA_4_ signalling. **(a)** SSeV infection assay showing the DUF1874 gene SIRV1 gp29 can neutralise the type III-B system in *S. islandicus*. Replicative plasmids with (pOE-gp29) or without (pOE) gp29 were transformed into the *S. islandicus* RJW007Δtype I-A mutant, and the resulting strains were then challenged with SSeV. Plaques are only observed on the lawn of the strain expressing gp29. Data are representative of three biological replicates, **(b)** Schematic showing the recombinant *M. tuberculosis* type III-A CRISPR interference system established in *E. coli*. By swapping the native Csm6 ancillary nuclease for a Csx1 protein, the system can be converted from cA_6_ to cA_4_-mediated immunity. **(c)** Plasmid transformation assay using a plasmid with a match to a spacer in the CRISPR array. If the plasmid is successfully targeted by the CRISPR system, transformants will not be seen. Plasmids with or without the *duf1874* gene are targeted successfully when cAβ (Csm6) mediated antiviral signalling is active. In contrast, cells using a cA_4_-based (Csx1) system only prevent transformation when the DUF1874 protein is not present, showing that the protein is effective in neutralising cA_4_-based CRISPR interference. The controls lack cOA-dependent ribonucleases.

To confirm the specificity of the anti-CRISPR function, we utilised a recently developed recombinant type III CRISPR system from *Mycobacterium tuberculosis*, which we have established in *E. coli*. The system allows the cOA effector protein to be swapped to provide effective immunity against mobile genetic elements based on either cA_6_ or cA_4_ signalling^10^. We set up an assay where cells capable of cA_4_ or cA_6_-based immunity were transformed with a plasmid that could be targeted for interference due to a match in the tetracycline resistance gene in the plasmid to a spacer in the CRISPR array (Figure 1b). Efficient interference was observed for either strain in the absence of the *duf1874* gene from bacteriophage THSA-485A (Figure 1c). However, the presence of the *duf1874* gene on the plasmid abrogated immunity for cA_4_, but not for cA_6_-mediated CRISPR defence. Thus, DUF1874 acts as an anti-CRISPR against cA_4_-mediated type III CRISPR defence, explaining the previous observations with archaeal viruses^24,25^. Hereafter we propose the collective name AcrIII-1 for this family, as it is the first Acr described for type III systems and is not specific for an individual subtype of the type III effectors^28^.

To explore the mechanism of the AcrIII-1 family, we cloned and expressed two family members in *E. coli:* the SIRV1 gp29 protein and the YddF protein encoded by an integrative and conjugative element ICEBs1 from *B. subtilis*^29^. Both proteins possess a potent ring nuclease activity, rapidly degrading cA_4_ to generate linear di-adenylate (ApA>P) with a cyclic 2’,3’ phosphate (Figure 2 and extended data figure 3). With a catalytic rate exceeding 5 min^−1^, the anti-CRISPR enzyme is at least 60-fold more active than the cellular ring nuclease Crn1 from *S. solfataricus*. We showed previously that the type III-D CRISPR effector of *S. solfataricus* generates cA_4_ in proportion to the amount of cognate target RNA present^16^. By varying target RNA input and following cA_4_ levels and Csx1 activity, we compared the abilities of Crn1 and AcrIII-1 to destroy the signalling molecule and deactivate the ancillary defence nuclease Csx1. In keeping with its low turnover number, the Crn1 enzyme was effective at degrading cA_4_ and thus deactivating Csx1 only at the lowest levels of target RNA (Figure 3a). In contrast, AcrIII-1 degraded cA_4_ completely at the highest target RNA concentration examined, preventing Csx1 activation. To provide a more rigorous test, we examined the ability of both enzymes to prevent Csx1 activation over a range of cA_4_ concentrations spanning four orders of magnitude (Figure 3b). Crn1 (2 *μ*M) provided protection only up to 5 *μ*M cA_4_, but in contrast 2 *μ*M AcrIII-1 provided complete protection at the highest level of cA_4_ tested (500 *μ*M). Thus, AcrIII-1 has the potential to destroy large concentrations of the cA_4_ second messenger rapidly, negating the immune response.

**Figure 2.**
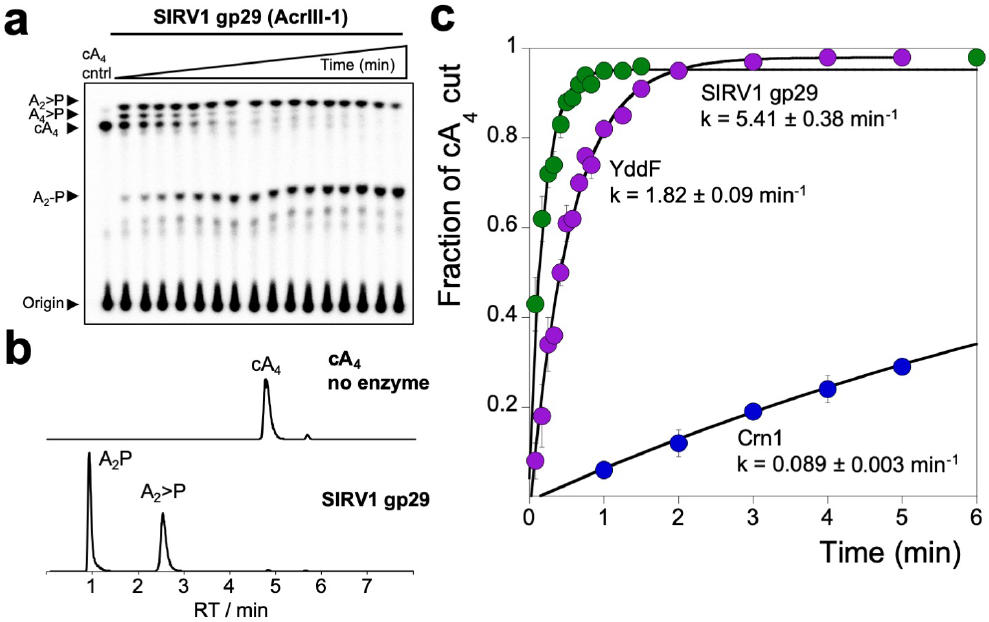
AcrIII-1 rapidly degrades cA_4_ to linear products. **(a)** Single-turnover kinetic analysis of cyclic tetra-adenylate (cA_4_, 200 nM) degradation by AcrIII-1 SIRV1 gp29 (4 *μ*M dimer, 50 °C), showing rapid generation of A_2_>P (di-adenylate with a 2’,3’-cyclic phosphate) via an A_4_>P intermediate visualised by phosphorimaging following thin-layer chromatography. A_2_>P is further hydrolysed to A2-P (di-adenylate with a 3’ phosphate). Each lane corresponds to reaction quenched at 5 second intervals up to 1 min, and then at 1.25, 1.5, 3, 6 and 12 min. Control reaction (cA_4_ cntrl) is cA_4_ incubated in the absence of protein for 12 min at 50 °C. **(b)** Liquid chromatography-high resolution mass spectrometry analysis confirms that AcrIII-1 SIRV1 gp29 converts cA_4_ (top panel) to A_2_>P within 2 min which is then converted to A2-P (bottom panel). (c) Kinetic comparison of cA_4_ degradation by the AcrIII-1 enzymes SIRV1 gp29 (4 *μ*M dimer, 50 °C) and YddF (8 *μ*M dimer, 37 °C) with the Crn1 enzyme Sso2081 (4 *μ*M dimer, 50°C). Data were quantified and plotted to show the fraction of CA_4_ cut over time and fitted to an exponential equation, as previously described^8^. For YddF and Crn1 the data shown is the average of three independent experiments and for SIRV1 gp29 the data is the average of two biological replicates comprising three independent replicates for each. The error indicated is the standard deviation of the mean.

**Figure 3.**
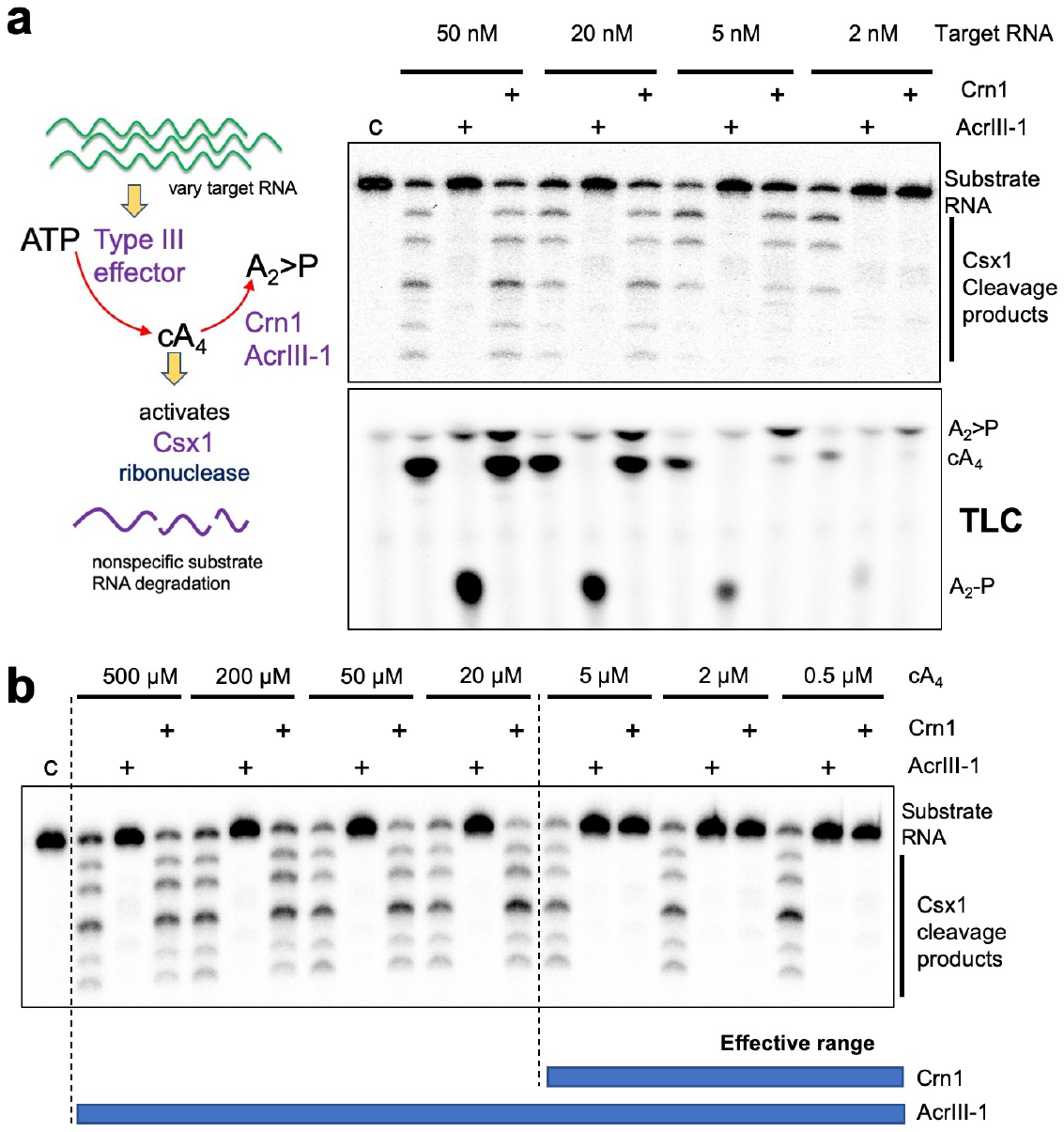
AcrIII-1 neutralises cA_4_-activated CRISPR defence enzyme Csx1. **(a)** Top panel is a phosphor image of denaturing PAGE visualising activation of Csx1 (0.5 μM dimer) in a coupled assay containing type III Csm complex when activated with indicated amounts of (unlabeled) target RNA to initiate cA_4_ synthesis. Each set of three lanes after the control (c) reaction with Csx1 and substrate RNA alone, is first in the absence and then with AcrIII-1 SIRV1 gp29 (2 μM dimer) or Crn1 Sso2081 (2 μM dimer), respectively. Whereas AcrIII-1 degraded all cA_4_ generated with up to 50 nM RNA target, the Crn1 enzyme deactivated Csx1 only when less than 5 nM RNA target was used to initiate cA_4_ synthesis. The lower panel is a phosphorimage of TLC with reactions as above but visualises cA_4_ production, by α-ATP incorporation, and degradation, in the presence of indicated amounts of RNA target and absence or presence of either AcrIII-1 or Crn1, and is representative of the results of three independent experiments. Csx1 deactivation correlates with complete cA_4_ degradation. **(b)** Denaturing PAGE showing activation of Csx1 (0.5 μM dimer) by indicated amounts (500-0.5 μM) of HPLC-purified cA_4_ and its subsequent deactivation when either AcrIII-1 or Crn1 (2 μM dimer) was present to degrade cA_4_. The AcrIII-1 enzyme degraded 100-fold more cA_4_ than Crn1. Control reaction (c) shows RNA incubated with Csx1 in the absence of cA_4_. The image is representative of the results of three independent experiments.

Several apo-structures for AcrIII-1 family members are available, revealing a dimeric protein of novel fold^22,23,30^. Importantly, this structure is completely unrelated to the CRISPR associated Rossman fold (CARF) domain, which is the only protein family thus far known to bind cOA^n^. To elucidate the mechanism of cA_4_ binding and cleavage by AcrIII-1, we co-crystallised an inactive variant (H47A) of SIRV1 gp29 with cA_4_ and solved the structure to 1.55 Å resolution (Figure 4). The complex reveals a molecule of cA_4_ bound at the dimer interface. Comparison of the cA_4_-bound and apo structures reveals a significant movement of a loop, comprising residues 82-94, and subsequent α-helix, to bury cA_4_ within the dimer. These loops adopt variable or unstructured conformations in the various apo protein structures^22,23,30^. Once bound, the ligand is completely enclosed by the protein – a considerable accomplishment when one considers the large size of the ligand and the small size of the protein (Figure 4b). By superimposing the CA_4_ ligand on the apo-protein structure, it becomes apparent that the binding site is largely pre-formed, with the exception of the mobile loops that form the lid (Figure 4c). The overall change is like two cupped hands catching a ball, with the loops (fingers) subsequently closing around it.

**Figure 4.**
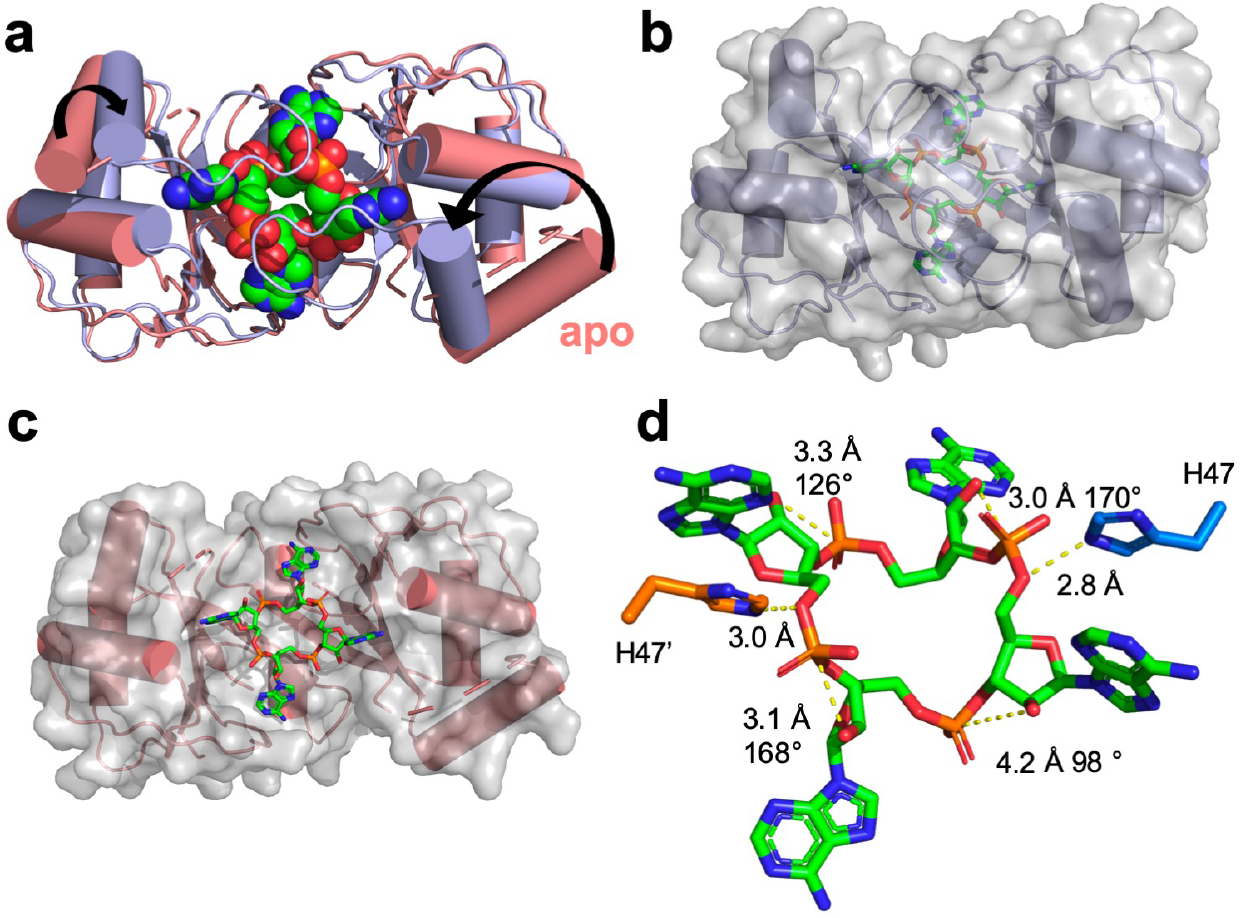
Structure of AcrIII-1 bound to cA_4_. **(a)** Superimposition of the apo SIRV1 gp29 structure (salmon) and in complex with cA_4_ (blue), highlighting the movement of the loop and α-helix upon cA_4_ binding. cA_4_ is shown in green spheres. **(b)** Surface representation of the structure of SIRV1 gp29 (blue) in complex with cA_4_ (green), emphasising the complete burial of the ligand. **(c)** Surface representation of the apo structure of SIRV1 gp29 (salmon) with cA_4_ (green) in the position observed in the complex structure, indicating that the binding site is pre-formed. **(d)** Structure of cA_4_ bound to SIRV1 gp29. The two active site histidine residues (modelled based on the position of the alanine side chain in the H47A variant crystallised with cA_4_; coloured to represent residues from different monomers) are in suitable positions to act as the general acid, protonating the oxyanion leaving group. The corresponding ribose sugars have 2’-hydroxyl groups suitably positioned for in-line nucleophilic attack on the phosphodiester bond.

The cA_4_ molecule makes symmetrical interactions with each monomer of AcrIII-1 (Extended data figure 4). Upon binding of cA_4_, arginine R85 on the loop from one monomer interacts with the distant half of the cA_4_ molecule and appears to ‘lock’ the closed dimer. Other important interactions are made with main chain L92,I69, and N8, and side chains R66, N8, Q81, S11, T50, S49, and N13, most of which are semi or fully conserved (Extended data figure 1 & 5), suggesting they have important roles in cA_4_ binding and / or catalysis in this whole family of enzymes.

At two positions, on opposite sides of the ring, the 2’-hydroxyl of the ribose is positioned correctly for in-line attack on the phosphodiester bond, consistent with the observed bilateral cleavage (Figure 4d). The catalytic power of the AcrIII-1 family likely derives from active site residues that position the 2’-hydroxyl group for in-line nucleophilic attack, stabilise the transition state and protonate the oxyanion leaving group^31^. For the AcrIII-1 family, the absolutely conserved residue His-47 is suitably positioned to act as a general acid and fulfil the latter role (Figure 4d). To test this hypothesis, we assayed variant H47A of AcrIII-1. The variant enzyme suffered a >2500-fold decrease in catalytic power, which could be partially reversed by chemical rescue with 500 mM imidazole in the reaction buffer (Extended data figure 6).

By targeting a key signalling molecule, a single AcrIII-1 enzyme should have broad utility in the inhibition of endogenous type III CRISPR systems in any species. The only constraint is the specificity for the cA_4_ signalling molecule. Of the CRISPR ancillary nucleases studied to date, most are activated by cA_4_; activation by cA_6_ appears to be limited to certain bacterial phyla including the Firmicutes and Actinobacteria^10^. By targeting the cA_4_ activator, AcrIII-1 will neutralize a slew of defence enzymes simultaneously. For example, *Thermus thermophilus* has two Csm6 enzymes^32,33^ and a recently described DNA nuclease, Can1^34^. All are activated by cA_4_ and would thus be deactivated by AcrIII-1. Recently, two other Acr proteins with enzymatic functions have been described: AcrVA1 which catalyses crRNA cleavage of Cas12a^35^ and AcrVA5, which acetylates the PAM-sensing site of Cas12a^36^. Both, however, target a protein (or protein:nucleic acid complex), implying a requirement for specific interactions that could be evaded by sequence variation.

The gene encoding AcrIII-1 is most prominent in the archaeal viruses, where it is found in representatives of at least five distinct viral families, making it one of the most widely conserved of all archaeal virus proteins^30^ (Extended data figure 1). Within archaeal genomes, homologues are typically adjacent to ORFs from mobile genetic elements rather than CRISPR loci, a good example being the STIV virus integrated into *S. acidocaldarius* genomes^37^. The distribution of AcrIII-1 in archaea is sporadic but covers most of the lineages, including crenarchaea, euryarchaea, thaumarchaea, Asgard archaea and others. AcrIII-1 is also present in several bacteriophages of the order *Caudovirales*, and there are many instances of *acrIII-1* genes in sequenced bacterial genomes, with homologues found in the Firmicutes, cyanobacteria, proteobacteria, actinobacteria and many more. Maximum likelihood phylogenetic analysis of the AcrIII-1 proteins suggests multiple horizontal gene transfers between unrelated viruses as well as between bacteria and archaea (Extended data figure 7). Sometimes the *acrIII-1* gene is clearly part of an integrated mobile genetic element as is the case for the *yddF* gene in *B. subtilis*^30^. However, in other species (n=49) the gene is associated with cellular type III CRISPR systems (Extended data figures 7–9). In *Marinitoga piezophilia*, AcrIII-1 is fused to a cOA-activated HEPN ribonuclease of the Csx1 family. Since both active sites are conserved, this fusion protein may have cA_4_ activated ribonuclease activity coupled with a cA_4_ degradative ring nuclease, thus providing an explicit linkage between the AcrIII-1 family and the type III CRISPR system. In this context the enzyme is likely acting as a host-encoded ring nuclease, like Crn1, rather than an Acr. We therefore propose the family name of Crn2 (CRISPR associated ring nuclease 2) to cover DUF1874 family members that are associated with type III CRISPR systems.

AcrIII-1 is the first Acr have functional roles in both viruses and cells^38^. It remains to be determined whether the *acrIII-1* gene arose in viruses and was appropriated by cellular type III systems or *vice versa*. However, the extremely broad distribution of *acrIII-1* and limited distribution of *crn2* suggests the former. Adoption of an anti-CRISPR protein as a component of a cellular CRISPR defence system seems counter-intuitive. However, the enzyme could be harnessed for a role in defence by either “detuning” its activity to make it a slower enzyme, or by putting it under tight transcriptional control so that it is expressed at very low levels. The unprecedented wide occurrence of this Acr across many archaeal and bacterial virus families reflects the fact that this enzyme degrades a key signalling molecule to subvert cellular immunity. This makes it very hard for cells to evolve counter-resistance, other than by switching to a different signalling molecule. The recent discovery of multiple cyclic nucleotide signalling systems in bacteria may thus reflect the ongoing conflict between cells and viruses.

## Acknowledgements

This work was supported by grants from the Biotechnology and Biological Sciences Research Council (REF: BB/S000313/1 to MFW and REF: BB/R008035/1 to TMG) and by a NASA Exobiology and Evolutionary Biology grant (NNX14AK23G to RJW). We thank Jesse Black and Maria Alejandra-Bautista for isolating and characterizing the SSeV virus, and thank Rebecca Wipfler and Wenlong Zhu for technical assistance.

## Author contributions

J.S.A. designed experiments and carried out the enzyme assays and analysis; S.M. carried out the structural biology; C.Z. constructed the *S. islandicus* strains and performed the virus infection assays; S.Grü. carried out the plasmid transformation assays and mass spectrometry; S.Gra. generated expression plasmids and purified proteins; M.R. contributed to the conception of the project; T.M.G., R.J.W. and M.F.W. oversaw the work, analysed the data and wrote the manuscript. All authors contributed to data analysis and writing.

## Correspondence

should be addressed to M.F.W. or T.M.G.

## Methods

### Construction of *S. islandicus* strains

The type I-A CRISPR defence module including seven genes i.e. *cas3b, csa5, cas7, cas5, cas3’, cas3’*, and *casX^39^* was in-frame deleted from the genetic host *S. islandicus* RJW007, derived from wild-type strain *S. islandicus* M.16.4 carrying a double *pyrEF* and *argD* deletion^26^, by employing Plasmid Integration and Segregation knockout strategy^40^. The type I-A deletion mutants were confirmed by PCR using primers that bind outside of the homologous flanking arms.

Synthesised SIRV1 *gp29* gene was purchased from IDT as a g-block and was cloned into a *Sulfolobus-E.coli* shuttle vector pSeSd-SsoargD^26^ (referred to as pOE hereafter) at the *NdeI* and NotI sites, generating the gp29 expression plasmid pOE-gp29 in which the *gp29* gene was placed under the control of the arabinose promoter. The pOE-gp29 and pOE plasmids were then transformed into competent cells of the Δtype I-A mutant via electroporation as described previously^26^, generating strains expressing and not expressing SIRV1 gp29, respectively.

### Viral quantification

To calculate the titer of SSeV, 100 μL diluted virus (10^−5^, 10^−6^, and 10^−7^) was co-incubated with 500 *μ*L *S. islandicus* Y08.82.36 host^27^ (10-fold concentrated) without shaking at 76-78 °C for 30 min. Afterwards, the virus-infected cells were transferred into a glass test tube containing 5 mL of pre-warmed Sucrose-Yeast (SY) and 0.8% gelrite mixture, and plated onto SY plates. The plates were put into a plastic bag, and incubated for two days at 76-78 °C. Plaques were counted in plates with proper virus dilutions, and the titer of SSeV were determined as 4.96×10^8^ plaque forming units (PFU) per ml.

### Infection by virus SSeV of *S. islandicus* M.16.4 with a type III CRISPR defence

The SSeV infection assay was carried out according to the procedure as described previously^41^ with minor modifications. In brief, approximately 6 × 10^8^ cells of RJW007Δtype I-A/pOE and RJW007Δtype I-A/pOE-gp29 taken from the exponential stage were spun down at 4000 rpm × 12 min, and resuspended in 1mL of Arabinose-Tryptone (AT) medium. The resuspensions were then co-incubated with 20 mL of fresh AT medium or SSeV supernatant at different dilutions (10^0^, 10^−1^, 10^−2^, 10^−3^, 10^−4^, 10^−5^, and 10^−6^) in a Falcon tube at 76-78 °C for 1 h without shaking. The SSeV-infected cells were washed twice with 10 mL of AT medium and resuspended into 500 *μ*L of AT medium. Afterwards, the concentrated SSeV-infected cells were mixed with 5 mL of top layer (2.5 mL of 2×Arabinose-Yeast medium+2.5 mL of 0.8% gelrite), and then plated onto the Arabinose-Yeast plates. PFU were counted after 4 days of incubation at 76-78 °C. Three independent experiments were performed.

### Cloning and purification

For cloning, synthetic genes (g-blocks) were purchased from Integrated DNA Technologies (IDT), Coralville, USA, and cloned into the vector pEhisV5spacerTev between the NcoI and *BamH*I sites^8^. Competent DH5a (*Escherichia coli*) cells were transformed with the construct and sequence integrity confirmed by sequencing (Eurofins Genomics). The plasmid was transformed into *Escherichia coli* C43 (DE3) cells for protein expression. Cloning of AcrIII-1 SIRV1 gp29, Crn1 Sso2081 and SsoCsx1 has been previously described^16,22^. For expression of SIRV1 gp29 and *Bacillus subtilis* YddF, 2 L of Luria-Broth culture was grown at 37 °C to an OD_600_ of 0.8 with shaking at 180 rpm. Protein expression was induced with 0.4 mM Isopropyl β-D-1-thiogalactopyranoside and cells were grown at 25 °C overnight before harvesting by centrifugation at 4000 rpm (Beckman Coulter Avanti JXN-26; JLA8.1 rotor) at 4 °C for 15 min.

For protein purification the cell pellet was resuspended in four volumes equivalent of lysis buffer containing 50 mM Tris-HCl 7.5, 0.5 M NaCl, 10 mM imidazole and 10% glycerol supplemented with EDTA-free protease inhibitor tablets (Roche; 1 tablet per 100 ml buffer) and lysozyme (1 mg/ml). Cells were lysed by sonicating six times 1 min with 1 min rest intervals on ice at 4 °C, and the lysate was ultracentrifuged at 40,000 rpm (70 Ti rotor) at 4 °C for 35 min. The lysate was then loaded onto a 5 ml HisTrap FF Crude column (GE Healthcare) equilibrated with wash buffer containing 50 mM Tris-HCl pH 7.5, 0.5 M NaCl, 30 mM imidazole and 10% glycerol. Unbound protein was washed away with 20 column volumes (CV) of wash buffer prior to elution of his-tagged protein using a linear gradient (holding at 10% for 3 CV, and 50% for 3 CV) of elution buffer containing 50 mM Tris-HCl pH 7.5, 0.5 M NaCl, 0.5 M imidazole and 10% glycerol. SDS-PAGE was carried out to identify fractions containing the protein of interest, and relevant fractions were pooled and concentrated using a 10 kDa molecular weight cut-off centrifugal concentrator (Merck). The his-tag was removed by incubating concentrated protein overnight with Tobacco Etch Virus (TEV) protease (1 mg per 10 mg protein) while dialysing in buffer containing 50 mM Tris-HCl pH 7.5, 0.5 M NaCl, 30 mM imidazole and 10% glycerol at room temperature. The protein with his-tag removed was isolated using a 5 ml HisTrapFF column, eluting the protein using 4 CV wash buffer. His-tag removed protein was further purified by size-exclusion chromatography (S200 16/60; GE Healthcare) in buffer containing 20 mM Tris-HCl pH 7.5, 0.125 M NaCl using an isocratic gradient. After SDS-PAGE, fractions containing protein of interest were concentrated and protein was aliquoted and stored at −80 °C. Variant enzymes were generated using the QuickChange Site-Directed Mutagenesis kit as per manufacturer’s instructions (Agilent technologies) and purified as for the wild-type proteins.

### Radiolabelled cA_4_ cleavage assays

Cyclic oligoadenylate (cOA) was generated by incubating 120 μg *Sulfolobus solfataricus (Sso*) type III-D (Csm) complex with 5 nM a-^32^P-ATP, 1 mM ATP, 120 nM A26 RNA target and 2 mM MgCl_2_ in Csx1 buffer containing 20 mM 2-(N-morpholino)ethanesulfonic acid (MES) pH 5.5, 100 mM K-glutamate, 1 mM DTT and 3 units SUPERase·In™ Inhibitor for 2 h at 70 °C in a 100 μl reaction volume. cOA was extracted by phenol-chloroform (Ambion) extraction followed by chloroform extraction (Sigma-Aldrich), and stored at −20 °C.

For single turnover kinetics experiments AcrIII-1 SIRV1 gp29 and variants (4 μM protein dimer) were assayed for radiolabelled cA_4_ degradation by incubating with 1 /400 diluted ^32^P-labelled SsoCsm cOA (~200 nM cA_4_; when generated in a 100 μl cOA synthesis reaction as described above) in Csx1 buffer supplemented with 1 mM EDTA at 50 °C. AcrIII-1 YddF (8 μM dimer) was incubated with cOA in buffer containing 20 mM MES pH 6.0, 100 mM NaCl, 1 mM DTT, 1 mM EDTA and 3 units SUPERase·In™ Inhibitor at 37 °C. For SIRV1 gp29 H47A chemical rescue, reactions were supplemented with 0.5 M imidazole. Two experimenters were involved in kinetic experiments involving 5 s time points. At desired time points, a 10 μl aliquot of the reaction was removed and quenched by adding to phenol-chloroform and vortexing. Subsequently, 5 μl of deproteinised reaction product was extracted into 5 μl 100% formamide xylene-cyanol loading dye if intended for denaturing polyacrylamide gel electrophoresis (PAGE), or products were further isolated by chloroform extraction if intended for thin-layer chromatography (TLC). A reaction incubating cOA in buffer without protein to the endpoint of each experiment was included as a negative control. All experiments were carried out in triplicate. For SIRV1 gp29 two biological samples were assayed in triplicate. cA_4_ degradation was visualised by phosphor imaging following denaturing PAGE (7M urea, 20% acrylamide, 1x TBE) or TLC.

For TLC, 1 μl of radiolabelled product was spotted 1 cm from the bottom of a 20 x 20 cm silica gel TLC plate (Supelco Sigma-Aldrich). The TLC plate was placed in a sealed glass chamber pre-warmed and humidified at 37 °C and containing 0.5 cm of a running buffer composed of 30% H_2_O, 70% ethanol and 0.2 M ammonium bicarbonate, pH 9.2. The temperature was lowered to 35 °C and the buffer was allowed to rise along the plate through capillary action until the migration front reached 17 cm. The plate was air dried and sample migration was visualised by phosphor imaging.

For kinetic analysis, cA_4_ cleavage was quantified using the Bio-Formats plugin^42^ of ImageJ as distributed in the Fiji package^43^ and fitted to a single exponential curve (y = m1 + m2*(1 – exp(-m3*x));m1=0.1;m2=1;m3=1;) using Kaleidagraph (Synergy Software), as described previously^44^. The cA_4_ cleavge rate by the H47A variant in the absence of imidazole was obtained by linear fit.

### Deactivation of HEPN nucleases by ring nucleases in coupled assays

In the absence and presence of Crn1 Sso2081 (2 μM dimer) or AcrIII-1 SIRV1 gp29 (2 μM dimer) 4 μg *S. solfataricus* Csm complex (~140 nM Csm carrying crRNA targeting A26 RNA target) was incubated with A26 RNA target (50, 20, 5, 2, or 0.5 nM) in buffer containing 20 mM MES pH 6.0, 100 mM NaCl, 1 mM DTT and 3 units SUPERase·In™ Inhibitor supplemented with 2 mM MgCl_2_ and 0.5 mM ATP at 70 °C for 60 min. 5’-end^32^P-labelled A1 RNA (AGGGUAUUAUUUGUUUGUUUCUUCUAAACUAUAAGCUAGUUCUGGAGA) and 0.5 μM dimer SsoCsx1 was added to the reaction at 60 min and the reaction was allowed to proceed for a further 60 min before quenching by adding phenol-chloroform. A1 RNA cleavage was visualised by phosphor imaging after denaturing PAGE. A control reaction incubating SsoCsx1 with A1 RNA in the absence of cOA was carried out to determine SsoCsx1 background activity. cA_4_ synthesis by Csm in response to A26 RNA target and subsequent cA_4_ degradation in the presence of Crn1 Sso2081 or AcrIII-1 SIRV1 gp29, was visualised by adding 5 nM a-^32^P-ATP with 0.5 mM ATP at the start of the reaction. Reactions were quenched at 60 min with phenol-chloroform and cA_4_ degradation products were visualised by phosphor imaging following TLC. A control reaction incubating Csm with ATP and a-^32^P-ATP in the absence of A26 RNA target was also carried out, quenching the reaction after 60 min.

cA_4_ degradation capacity of AcrIII-1 SIRV1 gp29 versus the Crn1 enzyme (Sso2081) was determined by incubating 2 μM dimer of each enzyme with 500-0.5 μM unlabelled cA_4_ (BIOLOG Life Science Institute, Bremen, Germany) in Csx1 buffer at 70 °C for 20 min before introducing SsoCsx1 (0.5 μM dimer) and ^32^P-labelled A1 RNA (50 nM). The reaction was left to proceed for a further 60 min at 70 °C before quenching by adding phenol-chloroform. Deproteinised products were separated by denaturing PAGE to visualise RNA degradation.

### Plasmid transformation assays

*Tsac 2833 mediated plasmid immunity from a reprogrammed type III system in E*. coliPlasmids pCsm1-5_ ΔCsm6 (containing the type III Csm interference genes *cas10, csm3, csm4, csm5* from *M. tuberculosis* and *csm2* from *M. canettii*), pCRISPR_TetR (containing *M. tuberculosis cas6* and tetracycline resistance gene-targeting CRISPR array), pRAT-Target (tetracycline-resistance, target plasmid) and *M. tuberculosis* (Mtb)Csm6/ *Thioalkalivibrio sulfidiphilus* (Tsu)Csx1 expression constructs have been described previously^10^. pRAT-Duet was constructed by replacing the pUC19 *lacZa* gene of pRAT-Target with the MCSs of pACYCDuet-1 by restriction digest (5’-NcoI, 3’-XhoI). The viral ring nuclease (*duf1784*) gene from *Thermoanaerobacterium* phage THSA_485A, tsac_2833, was PCR-amplified from its pEHisTEV expression construct and cloned into the 5’-NdeI, 3’-XhoI sites of MCS-2. The cOA-dependent nucleases (*mtb csm6, tsu csx1*) were cloned into the 5’-*Nco*I, 3’-*Sal*I sites of MCS-1 by restriction digest from their respective expression constructs. Each nuclease was cloned with and without the viral ring nuclease; pRAT-Duet without insert and pRAT-Duet containing only the viral ring nuclease were used as controls. The plasmid transformation assay was carried out essentially as described in^10^. *E. coli* C43 containing pCsm1-5_DCsm6 and pCRISPR_TetR were transformed by heat shock with 100 ng of pRAT-Duet target plasmid containing different combinations of cOA-dependent nuclease and viral ring nuclease. After outgrowth at 37 °C for 2 h, cells were collected and resuspended in 200 μl LB. A series of 10-fold dilutions was applied onto LB agar containing 100 μg ml^−1^ ampicillin and 50 μg ml^−1^ spectinomycin to determine the cell density of the recipient cells and onto LB agar additionally containing 25 μg ml^−1^ tetracycline, 0.2% (*w/v*) D-lactose and 0.2% (*w/v*) L-arabinose to determine the cell density of viable transformants. Plates were incubated at 37 °C for 16 – 18 h; further incubation was carried out at room temperature.

### Liquid chromatography high-resolution mass spectrometry

AcrIII-1 SIRV1 gp29 (40 μM dimer) was incubated with 400 μM cA_4_ in Csx1 buffer for 2 min at 70 °C and deproteinised by phenol-chloroform extraction followed by chloroform extraction. Liquid chromatography-high resolution mass spectrometry (LC-HRMS) analysis was performed on a Thermo Scientific™ Velos Pro instrument equipped with HESI source and Dionex UltiMate 3000 chromatography system. Compounds were separated on a Kinetex^®^ EVO C18 column (2.6 *μ*m, 2.1’ 50 mm, Phenomenex) using the following gradient of acetonitrile (B) against 20 mM ammonium bicarbonate (A): 0 – 2 min 2% B, 2 – 10 min 2 – 8% B, 10 – 11 min 8 – 95% B, 11 – 14 min 95% B, 14 – 15 min 95 – 2% B, 15 – 20 min 2% B at a flow rate of 300 *μ*l min^−1^ and column temperature of 40 °C. UV data were recorded at 254 nm. Mass data were acquired on the FT mass analyzer in negative ion mode with scan range *m/z* 150 – 1500 at a resolution of 30,000. Source voltage was set to 3.5 kV, capillary temperature was 350 °C, and source heater temperature was 250 °C. Data were analysed using Xcalibur™ (Thermo Scientific).

### Phylogenetic analysis

AcrIII-1 homologs were collected by using gp29 (NP_666617) of SIRV1 as a query and running two iterations (E=1e-05) of PSI-BLAST^45^ against the non-redundant protein database at NCBI. The sequences were aligned using PROMALS3D^46^. Redundant sequences (95% identity threshold) and sequences with mutated H47 active site residue were removed from the alignment. Poorly aligned (low information content) positions were removed using the gt 0.2 function of Trimal^47^. The final alignment contained 124 positions. The maximum likelihood phylogenetic tree was constructed using the PhyML program^48^ with the automatic selection of the best-fit substitution model for a given alignment. The best model identified by PhyML was LG +G+I. The branch support was assessed using aBayes implemented in PhyML. The tree was visualized using iTOL^49^.

### Crystallisation

AcrIII-1 H47A variant was concentrated to 10 mg ml^−1^, incubated at 293 K for 1 hour with a 1.2 M excess of cA_4_, and centrifuged at 13,000 rpm for 10 minutes prior to crystallization. Sitting drop vapor diffusion experiments were set up at the nanoliter scale using commercially available and in-house crystallization screens and incubated at 293 K. Crystals appeared in various conditions, but those used for data collection grew from 40% 2-methyl-2,4-pentanediol, 5% polyethylene glycol 8000 and 0.1 M sodium cacodylate, pH 6.5. Crystals were harvested and transferred briefly into cryoprotectant containing mother liquor with 20% glycerol immediately before cryo-cooling in liquid nitrogen. The H47A variant was used to avoid cleavage of the cA_4_ substrate during co-crystallisation. The position of the active site histidine was inferred from the structure of the apo-protein.

### Data collection and processing

X-ray data were collected from two crystals at 100 K, at a wavelength 0.9686 Å, on beamline I24 at Diamond Light Source, to 1.49 and 1.60 Å resolution. Both data sets were automatically processed using Xia2^50^, utilizing XDS and XSCALE^51^. The data were merged in Aimless^52^ and the overall resolution truncated to 1.55 Å. The data were phased by molecular replacement using Phaser^53^ with a monomer from PDB file 2X4I stripped of water molecules as the search model. Model refinement of AcrIII-1 was achieved by iterative cycles of REFMAC5^54^ in the CCP4 suite^55^ and manual manipulation in COOT^56^. Electron density for cA_4_ was clearly visible in the maximum likelihood/*σ_A_* weighted *F*_obs_ – *F*_calc_ electron density map at 3σ. The coordinates for cA_4_ were generated in ChemDraw (Perkin Elmer) and the library was generated using acedrg^57^, before fitting of the molecule in COOT. Model quality was monitored throughout using Molprobity^58^. Data and refinement statistics are shown in Table S1.

### Data availability statement

The structural coordinates and data have been deposited in the Protein Data Bank with deposition code 6SCF. Raw data is available for the plasmid immunity analysis presented in figure 1.

Raw data is available for the kinetic analysis presented in figure 2 and extended data figure 6.

## Extended dataset

**Extended data figure 1.**
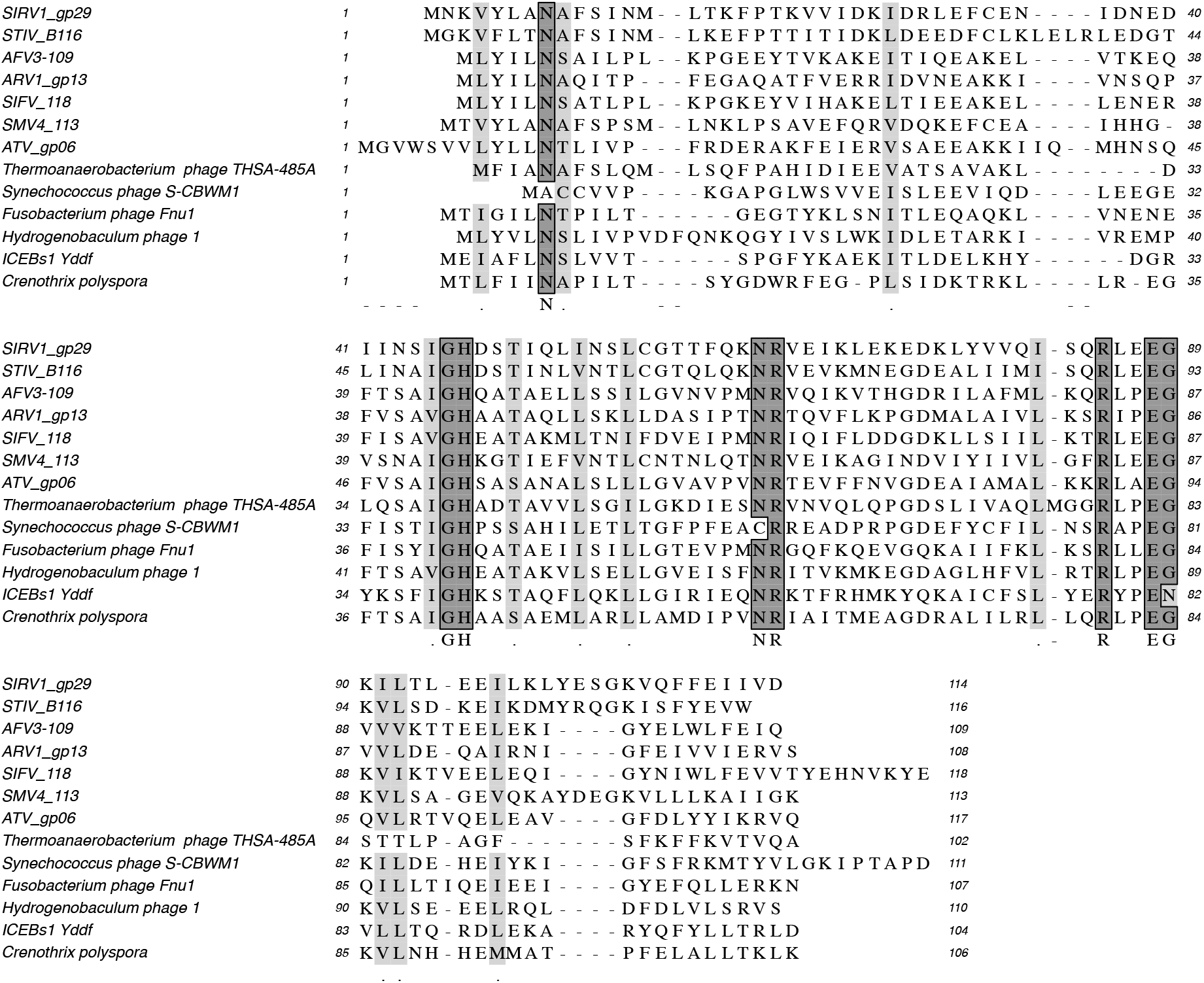
Multiple sequence alignment of DUF1874 family members. Includes the AcrIII-1 proteins from the archaeal viruses SIRV1, STIV, AFV3, ARV1, SIFV, SMV4 and ATV, the integrated conjugative element ICEBs1 protein YddF from *B. subtilis*, the bacteriophage proteins from *Thermoanaerobacterium* phageTHSA-485A, *Synechococcus* phage S-CBWM1, *Fusobacterium* phage Fnu1, *Hydrogenobaculum* phage 1 and the Crn2 protein from *Crenothrix polyspora*.

**Extended data figure 2.**
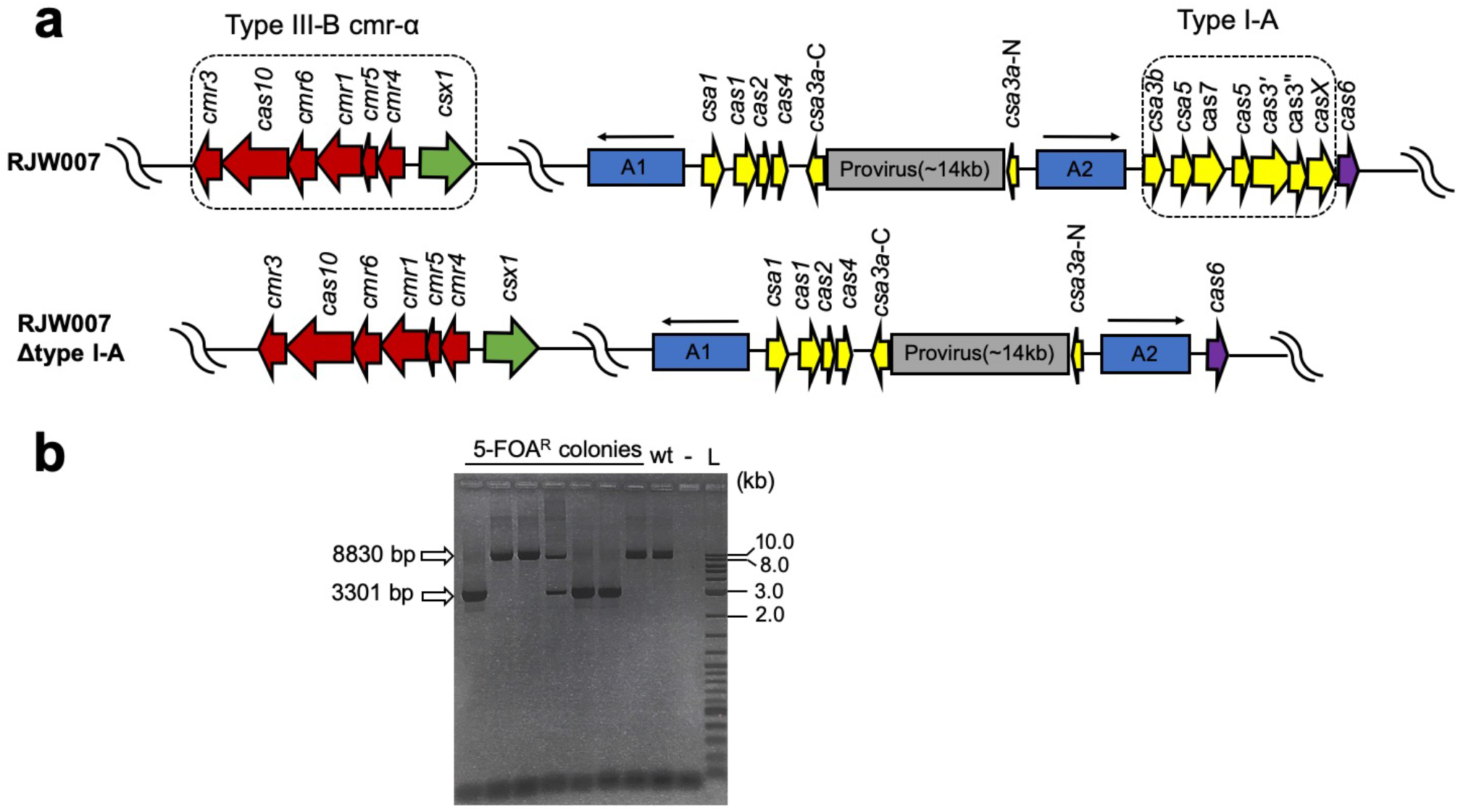
Construction of type I-A deletion mutant in the genetic host *S. islandicus* RJW007 (S. *islandicus* M.16.4Δ*pyrEF*Δ*argD*). **(a)** Genomic context of CRISPR system in the genetic host and the mutant strain. A1 and A2 denote two different CRISPR arrays, the orientation of which are indicated with arrows. **(b)** PCR verification of Δtype I-A mutants. A representative *Sulfolobus* transformant with type I-A knockout plasmid integrated was grown in Dextrin-Tryptone liquid medium, and the cell cultures were subsequently plated on Dextrin-Tryptone plates containing 5-FOA (5-fluoroorotic acid, 50 *μ*g/mg), uracil (20 *μ*g/ml), and agamatine (1 mg/ml). Seven randomly selected 5-FOA resistant (5-FOA^R^) colonies were screened using the primer set type IA-flankP-For/Rev that binds outside of the flanking homologous regions to confirm the type I-A deletion. A representative Δtype I-A mutant was further colony-purified for following experiment. The expected size of PCR product amplified from the genomic DNA of parental strain (referred to wt) and Δtype I-A mutant is 8830 bp and 3001 bp, respectively. −, negative control (water as DNA template for PCR). L, 2-log DNA ladder (NEB, USA).

**Extended data figure 3.**
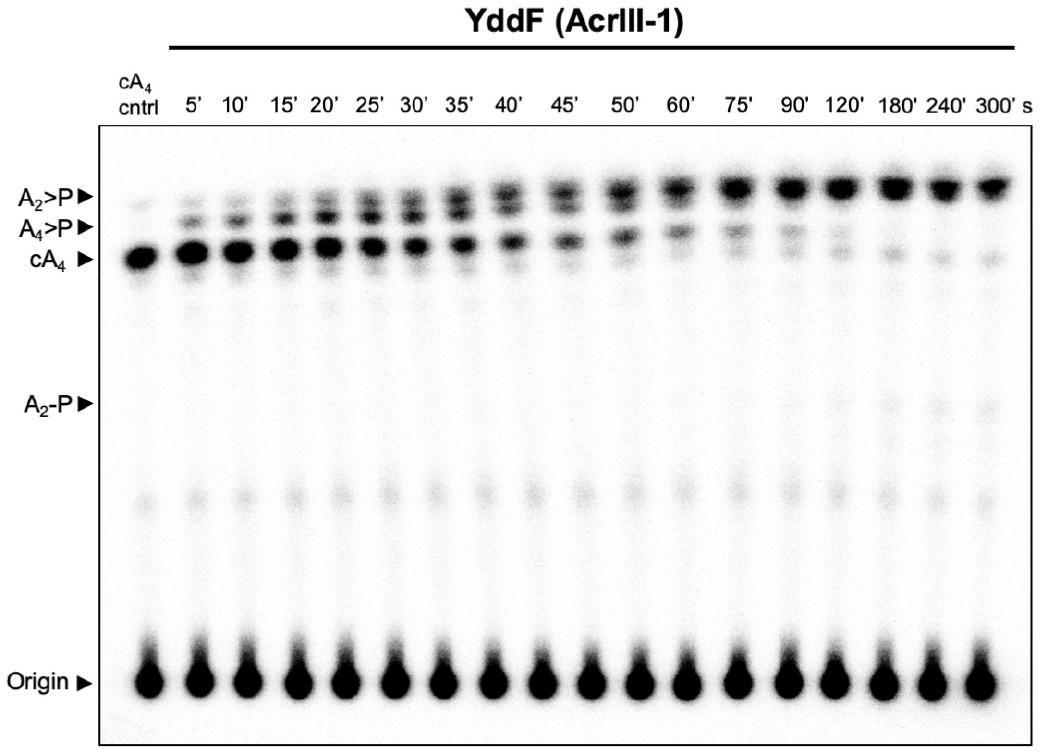
Single turnover kinetics of cA_4_ cleavage by *B. subtilis* YddF. Phosphorimage of thin-layer chromatography (TLC) visualising cA_4_ cleavage by YddF (8 μM dimer, 37°C) over a period of 5 min. The rate of cA_4_ cleavage to generate A_4_>P (intermediate product tetra-adenylate containing 2’,3’ phosphate and a 5’ hydroxyl moiety) and A_2_>P over time (as shown in figure 2c) was calculated by quantifying densiometric signals from the phosphorimage, which is representative of three independent experiments. Control reaction (cA_4_ cntrl) is cA_4_ incubated in the absence of protein for 5 min at 37°C.

**Extended data figure 4.**
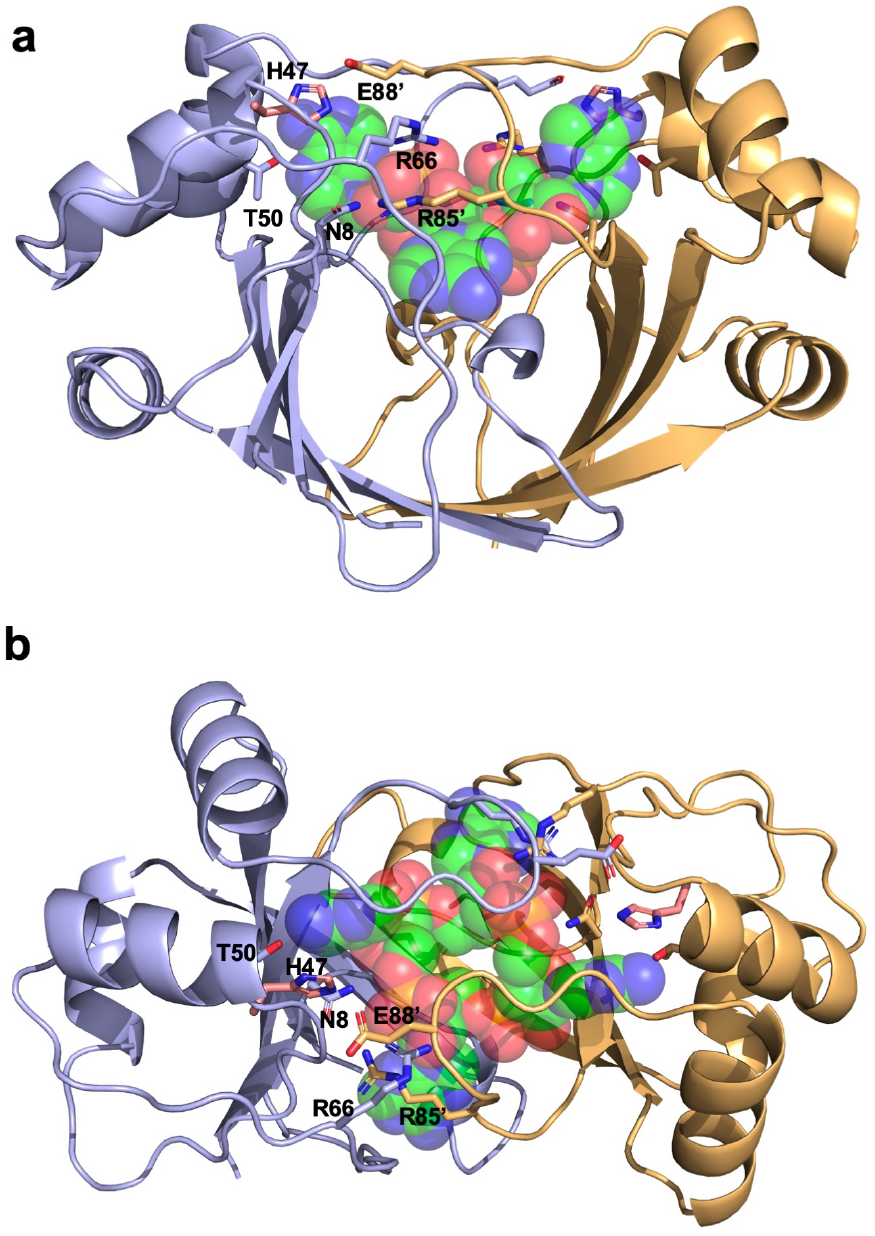
Structure of SIRV1 gp29 bound to cA_4_. Orthogonal views of SIRV1 gp29 dimer in complex with cA_4_. The monomers are coloured blue and orange, with catalytic residue H47 from the apo structure shown in salmon. cA_4_ is shown in green spheres. Conserved residues (Extended data figure 1) in the AcrIII-1 family are indicated and discussed in the text.

**Extended data figure 5.**
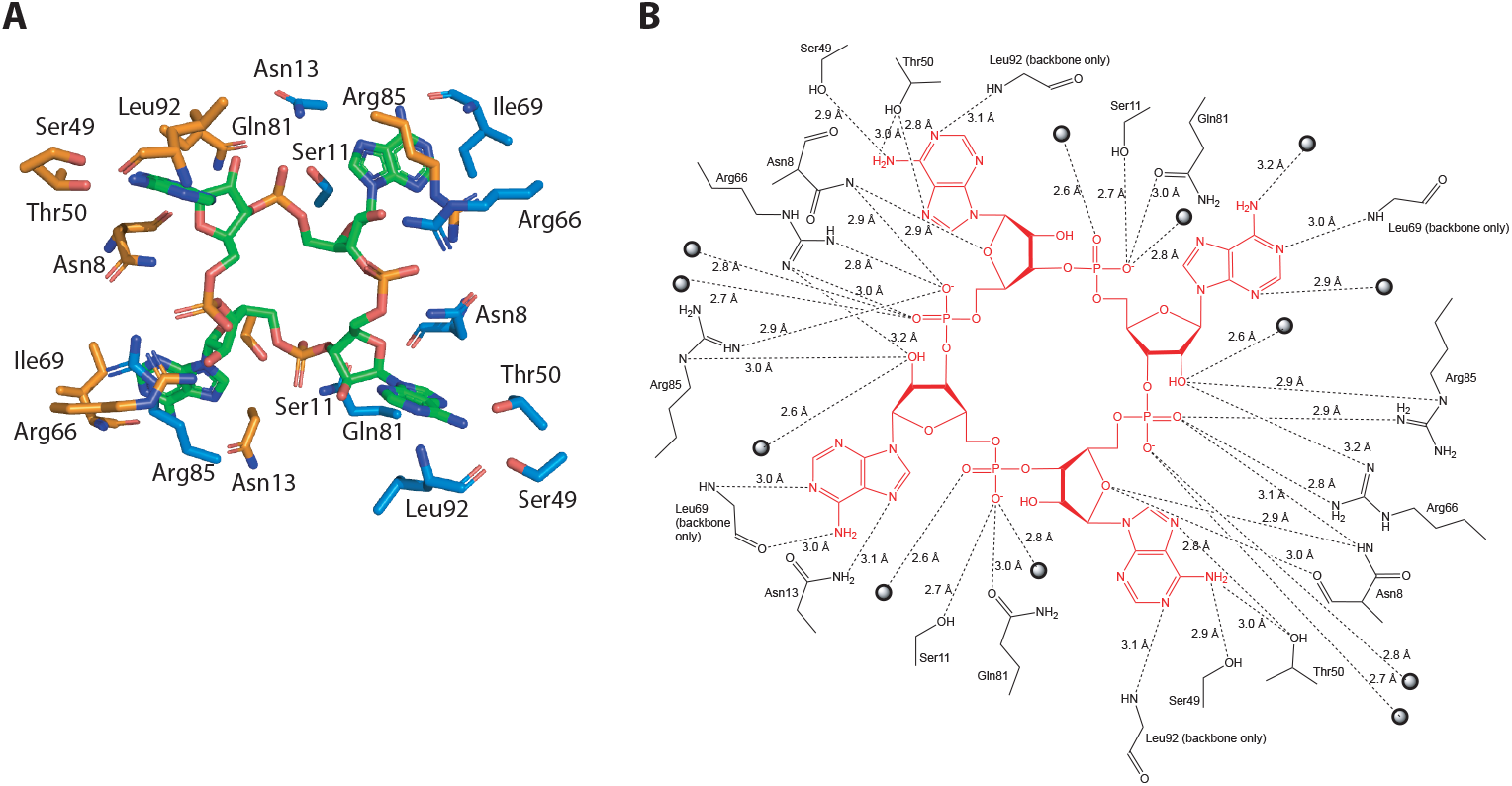
Interactions made in the active site of SIRV1 gp29 in complex with cA_4_. **(a)** Interactions between each monomer of the SIRV1 dimer (orange and blue), with cA_4_ shown in green. **(b)** Schematic showing the interaction. Dotted lines represent hydrogen bonds, with the distance annotated. Spheres represent water molecules.

**Extended data figure 6.**
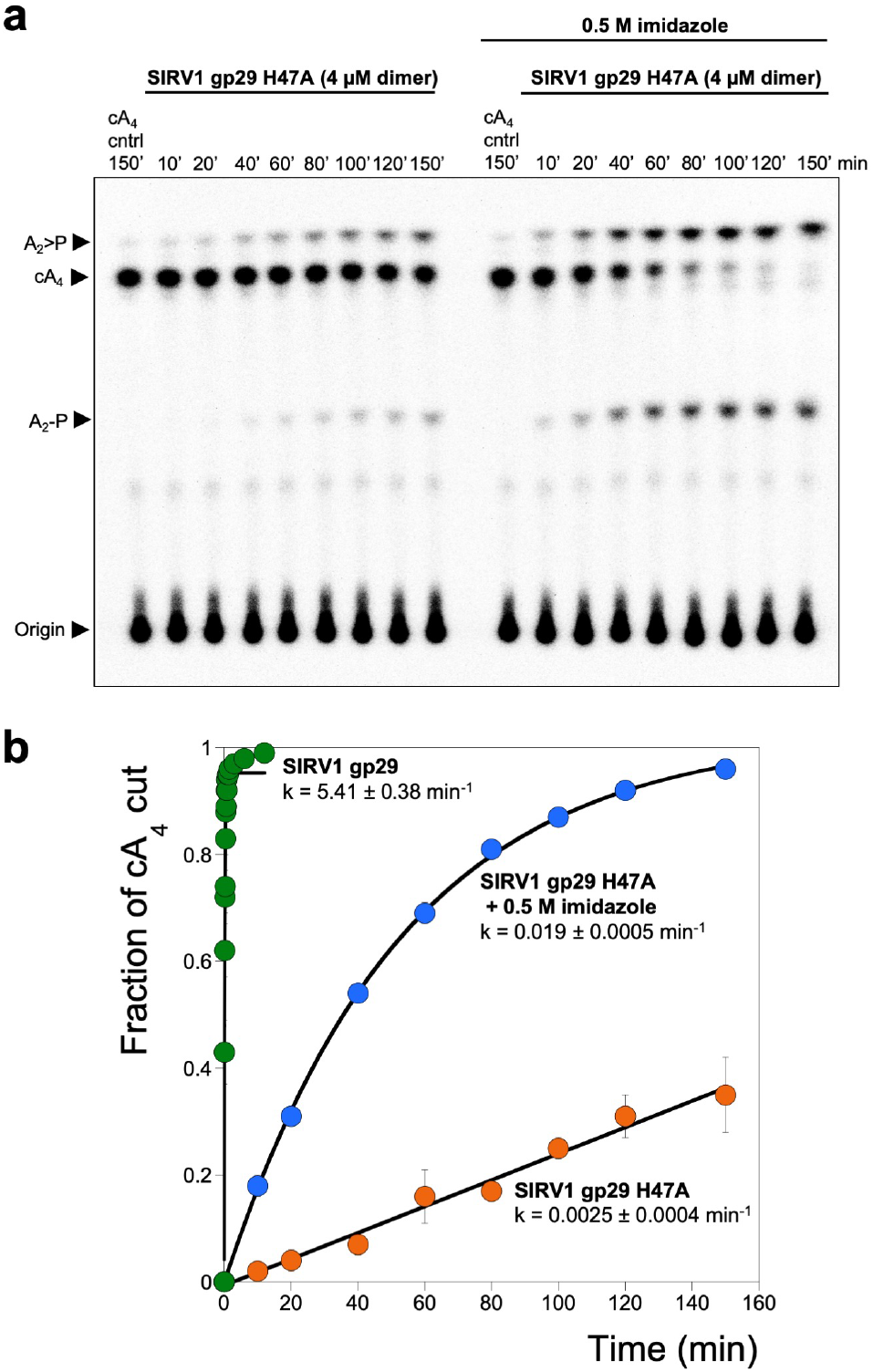
Single turnover cA_4_ cleavage by SIRV1 gp29 H47A and chemical rescue with imidazole. **(a)** Phosphorimage of TLC visualising cA_4_ cleavage by SIRV1 gp29 H47A (4 μM dimer, 50 °C) in the presence or absence of 500 mM imidazole, over time. The rate of cA_4_ cleavage to generate A_2_>P and A_2_-P was calculated by quantifying densiometric signals from the phosphorimage, which is representative of three independent experiments. **(b)** Plot comparing the single turnover rates of cA_4_ by SIRV1 gp29 and its H47A variant in the presence or absence of imidazole. cA_4_ cleavage by the H47A variant can be partially restored when the reaction is supplemented with 500 mM imidazole. Errors bars indicate the standard deviation of the mean.

**Extended data figure 7.**
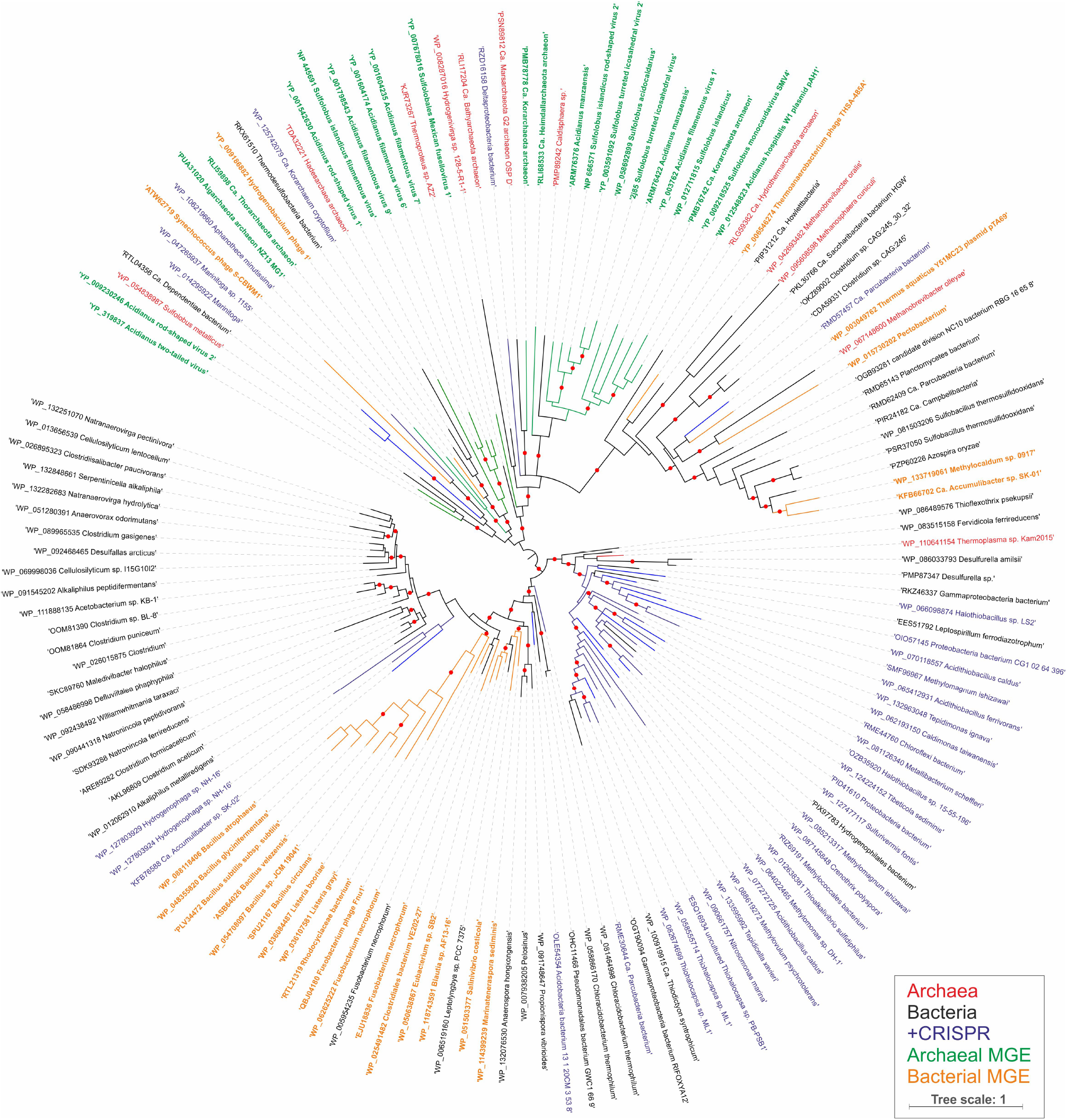
Maximum likelihood phylogeny of AcrIII-1 homologs. The maximum likelihood phylogenetic tree was constructed with the automatic selection of the best-fit substitution model for a given alignment (LG +G+I). Red circles indicate 95-100% branch support, as assessed using aBayes implemented in PhyML. The scale bar represents the number of substitutions per site. Branches and labels are colour coded: archaea, red; bacteria, black; bacteria and archaea in which AcrIII-1 homologs are associated with CRISPR loci, blue; archaeal viruses and plasmids, green; bacteriophages, orange.

**Extended data figure 8.**
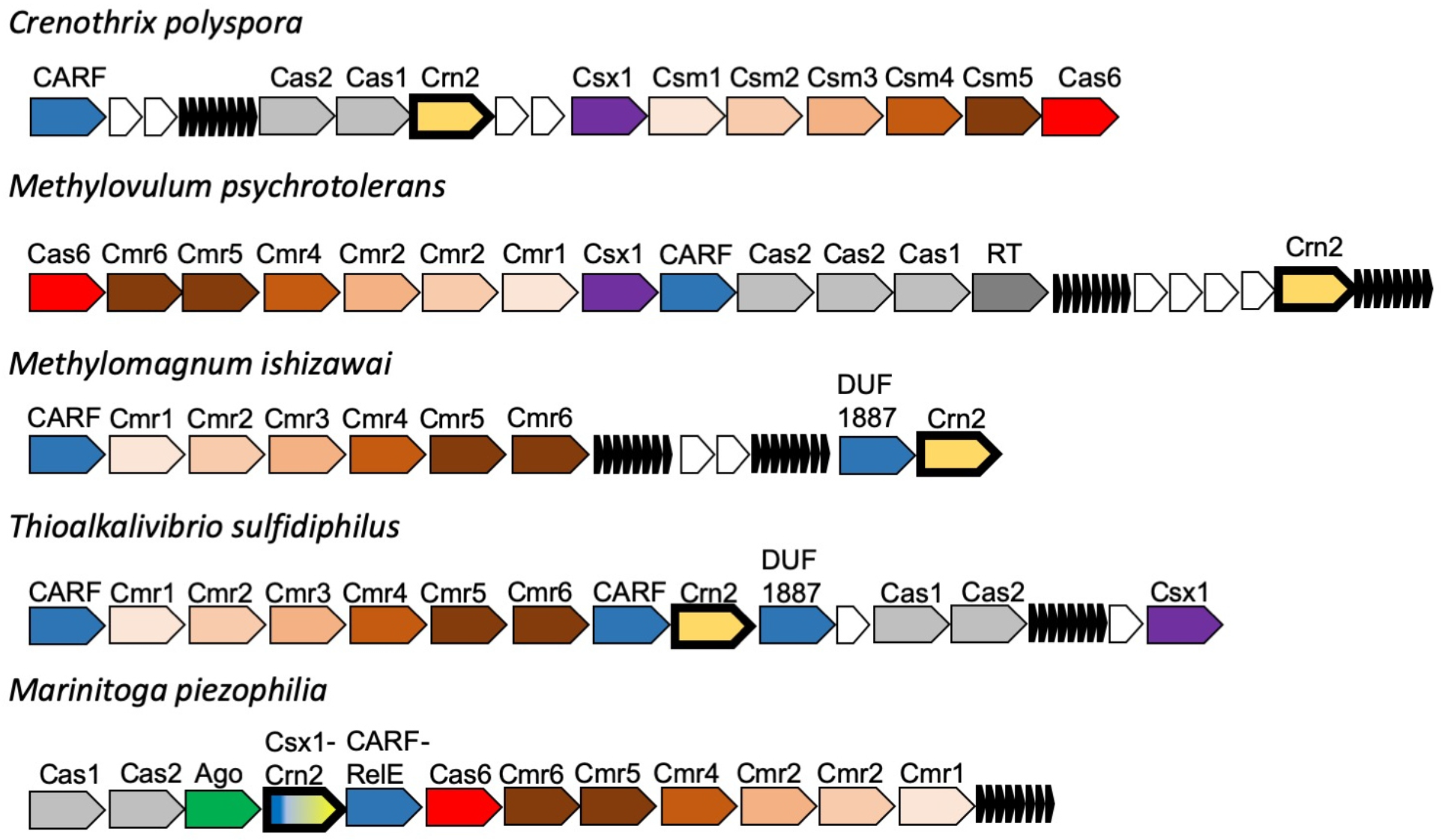
Genomic context of *crn2* genes in selected bacteria. Type III CRISPR loci in the bacterial species *Crenothrix polyspora, Methylovulum psychrotolerans, Methylomagnum ishizawai, Thioalkalivibrio sufidiphilus* and *Marinitoga piezophilia* are shown in cartoon form, with genes labelled and colour-coded. The *crn2* gene is shown in pale yellow with a bold outline; CRISPRs are indicated by small black arrows and unrelated/hypothetical genes shown as small white arrows. The size and orientation of genes is not reflected in the cartoon. Key to gene labels: RT, Reverse Transcriptase; Ago, Argonaute; CARF, CRISPR associated Rossman fold; DUF1887, predicted CARF nuclease; CARF-RelE, CARF domain fused to the RelE toxin.

**Extended data figure 9.**
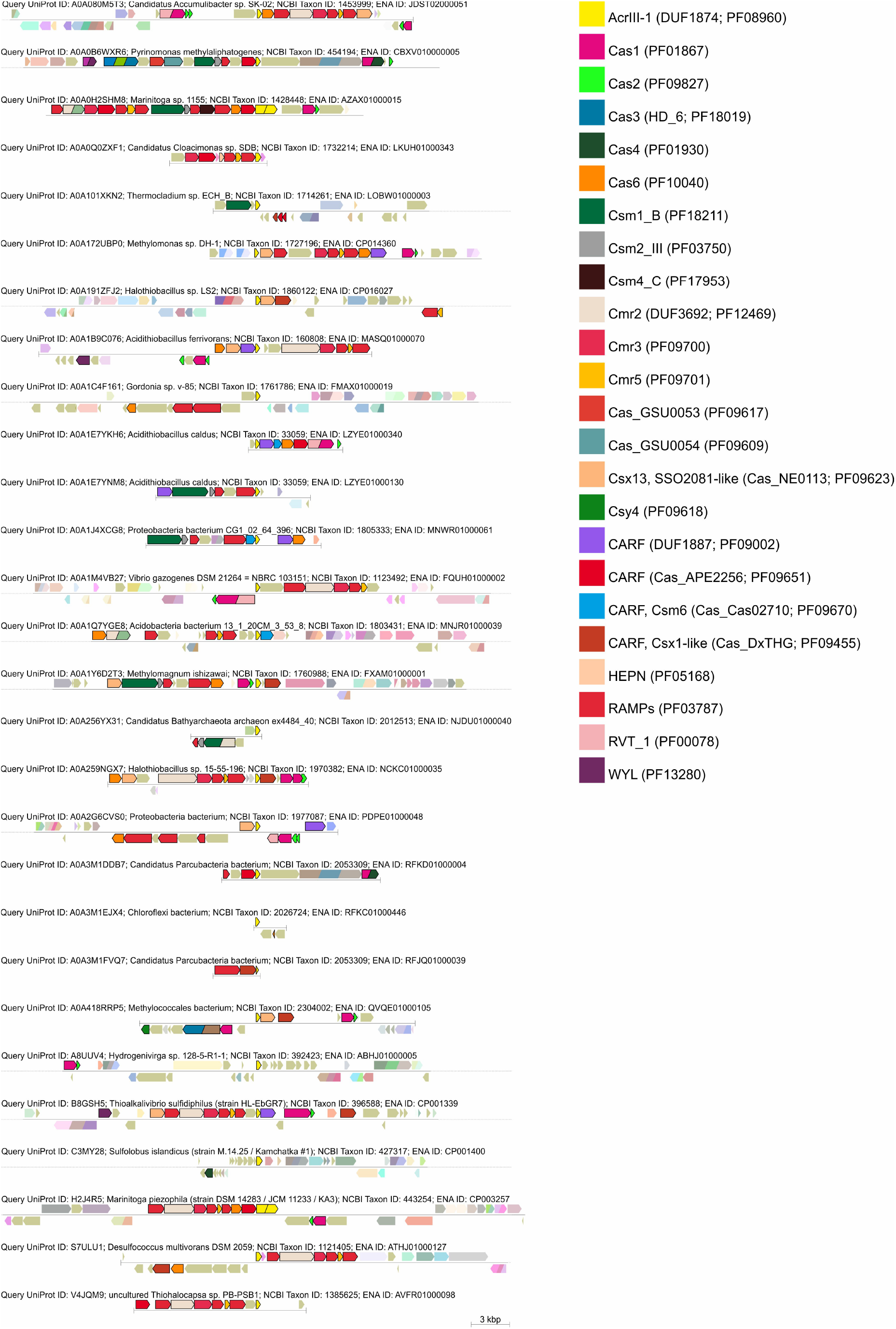
CRISPR-associated ACRIII-1 homologs. Genomic neighbourhoods were analysed using the Enzyme Function Initiative-Genome Neighbourhood Tool (EFI-GNT) against the Pfam profile database^59^. Gene annotations are colour coded and the key is provided on the right of the figure.

**Table S1:**
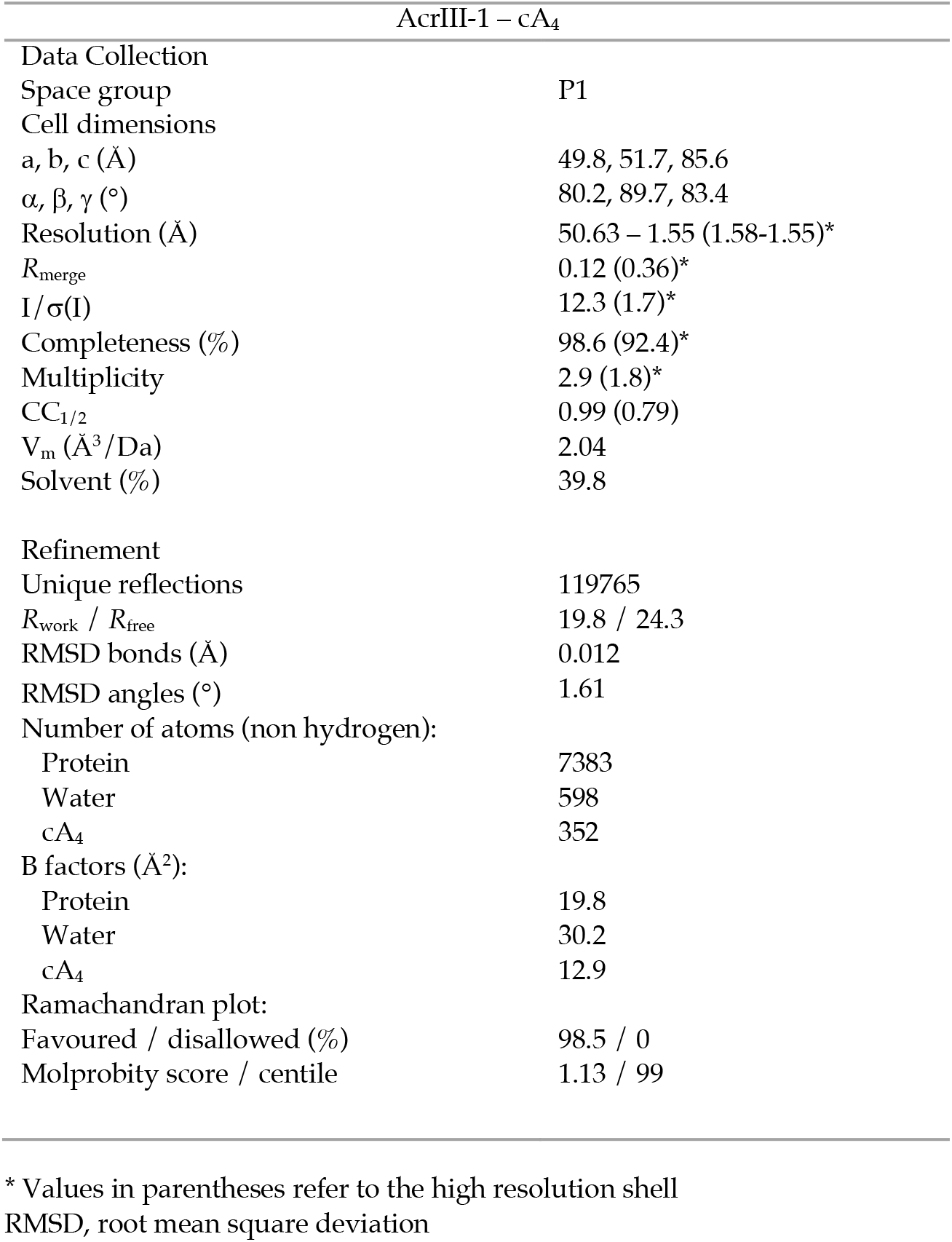
Data collection and refinement statistics for AcrIII-1 in complex with cA_4_

